# A Computational Synaptic Antibody Characterization and Screening Framework for Array Tomography

**DOI:** 10.1101/258756

**Authors:** Anish K. Simhal, Belvin Gong, James S. Trimmer, Richard J. Weinberg, Stephen J. Smith, Guillermo Sapiro, Kristina D. Micheva

## Abstract

Application-specific validation of antibodies is a critical prerequisite for their successful use. Here we introduce an automated framework for characterization and screening of antibodies against synaptic molecules for high-resolution immunofluorescence array tomography (AT). The proposed Synaptic Antibody Screening Tool (SACT), is designed to provide an automatic, robust, flexible, and efficient tool for antibody characterization at scale. By allowing the user to define the molecular composition and size of synapses expected to contain the antigen, the method detects and characterizes puncta and synapses, and outputs automatically computed characteristics such as synapse density and target specificity ratio, which reflect the sensitivity and specificity of immunolabeling with a given antibody. These measurements provide an objective way to characterize and compare the performance of different antibodies against the same target, and can be used to objectively select the antibodies best suited for AT and potentially for other immunolabeling applications.

## 1 INTRODUCTION

Antibodies are an indispensable tool for the modern biologist. Their high-affinity binding to specific target molecules makes it possible to detect, isolate, and manipulate the function of these molecules. A staggering number of antibodies are available to the research community, as are many options to make new antibodies. However, since antibodies are biological tools employed in complex systems, they can be very difficult to evaluate and to use in a predictable and reproducible way. A large volume of misleading or incorrect data has been published based on results from antibodies that did not perform as assumed (Anderson and Grant, 2006; Baker, 2015; Rhodes and Trimmer, 2006). Recognizing this problem, there has been substantial progress in optimizing antibody production (Nilsson et al., 2005; Gong et al., 2016). The importance of establishing reliable practices for antibody use is now widely accepted and many companies are adopting transparent practices for rigorous antibody validation (Fritschy et al., 1998; Uhlen et al., 2016). The performance of antibodies, however, is application-specific (Lorincz and Nusser, 2008), and the reliable performance of an antibody in one application does not guarantee its suitability for another application. For example, an antibody that yields a single band on an immunoblot analysis of a tissue homogenate may prove wholly unsatisfactory for immunohistochemistry on sections of fixed tissue. Moreover, the same antibody that yields a robust and specific signal in immunohistochemical labeling of tissue sections prepared under one set of conditions may yield a weak or noisy signal on comparable samples prepared under different conditions (Fritschy et al., 1998; Fukaya and Watanabe, 2000). Therefore, it is up to the individual users to validate antibodies for other applications and conditions. This task is especially crucial for applications whose chemistry differs substantially from standard immunoblots.

Array tomography (AT) is a technique which involves immunolabeling and imaging of serial arrays of ultrathin (*∼*70 nm) plastic-embedded tissue slices of aldehyde-fixed tissue (Micheva and Smith, 2007; Micheva et al., 2010). While embedding tissue in resin has multiple advantages, the embedding process requires tissue dehydration, infiltration in plastic resin, and subsequent resin polymerization, all of which can modify the protein structure and chemical state and have a major impact on its immunoreactivity. Therefore, identifying antibodies that yield robust and specific immunolabeling of target proteins in plastic sections is a daunting task. A particular topic of great biological interest is to better understand synapses in the mammalian brain, and finding antibodies that selectively label subcellular targets like synapses presents additional challenges, due to their small size, high density, and overall complexity.

The primary criterion for evaluating antibody performance for a given application is determining whether the labeling pattern is consistent with the known tissue characteristics of its target protein. For an antibody against a synaptic protein, the immunolabeling must be localized at synapses. Though conceptually straightforward, the practical evaluation of this criterion is difficult and often involves a number of subjective and time-consuming decisions. A synapse can be unambiguously identified via electron microscopy, but this approach is too time-consuming and expensive to be practical for large scale (*>*100 antibody clones) antibody screening tests. A more efficient strategy is to double label the same sample with another antibody already known to localize at synapses, and measure colocalization (Micheva et al., 2010; Weiler et al., 2014). While effective, this method presents a number of challenges. Synaptic proteins are typically expressed in high concentrations at synapses; however, these proteins are also present to various (and often unknown) degrees at other subcellular locations. Furthermore, synapses display a high level of protein diversity, so many synapses may completely lack a particular synaptic protein. Adding to the uncertainty, other sources of fluorescence can confuse the interpretation of the images. These sources of ‘noise’ include signal arising from autofluorescent tissue constituents such as lipofuscin granules, blood cells, contamination, and defects such as tissue folds created during sample processing as diagrammed in Figure 1. The trained eye of an expert can usually discern the different fluorescence sources, and pick out the specific immunolabel, however this is a subjective and non-quantitative assessment. Furthermore, it can be extremely time consuming, especially when examining a a large number of antibodies against the same antigen, as may be required during antibody production (Gong et al., 2016).

**Figure 1:**
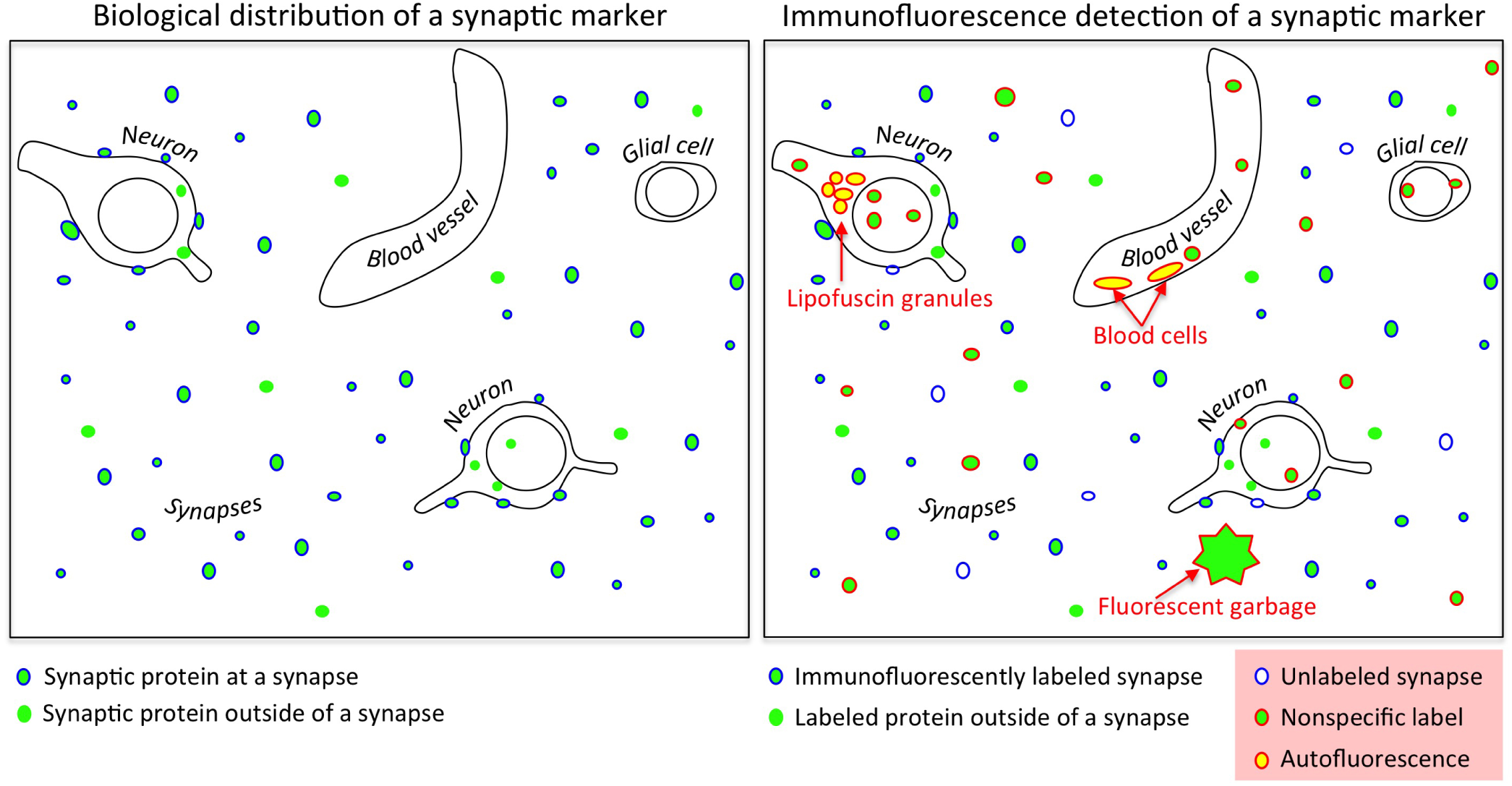
Challenges in evaluating synaptic antibodies. *Left.* Synaptic proteins are found not only at synapses, but also outside of synapses at sites of synthesis and transport; many synaptic proteins also lie in other subcellular compartments reflecting other functions (such as transcriptional regulators). *Right.* Immunofluorescence detection of synaptic proteins is confounded by nonspecific binding of antibodies, low efficiency of target protein detection, fluorescent contaminants, and tissue autofluorescence (e.g., lipofuscin granules and blood cells).

Accordingly, there is an urgent need for an efficient and robust framework for evaluation of synaptic antibodies. Here, we introduce the Synaptic Antibody Characterization Tool (SACT), which provides automatic and quantitative measurements of the intensity and specificity of immunolabel and enables the objective characterization and comparison of multiple synaptic antibodies for AT at scale.

## 2 METHODS

### 2.1 Overview

The proposed Synaptic Antibody Characterization Tool (SACT) was developed for the quantitative assessment of antibodies against synaptic targets used for AT. It computes key characteristics and measures of antibody performance, such as density, size and size variability of immunofluorescent puncta, as well as density of synapses detected by the antibody and the “target specificity ratio” (ratio of synapses detected to puncta detected). These measures provide unbiased quantitative information to help evaluate the specificity and sensitivity of immunolabeling obtained with a given antibody. The approach is outlined in Figure 2. The proposed SACT combines automatic synapse detection from (Simhal et al., 2017) with a novel puncta detection and characterization computational tool. We should note that SACT is a framework; additional measurements can be added and adapted as needed depending on the desired antibody characterization features.

**Figure 2:**
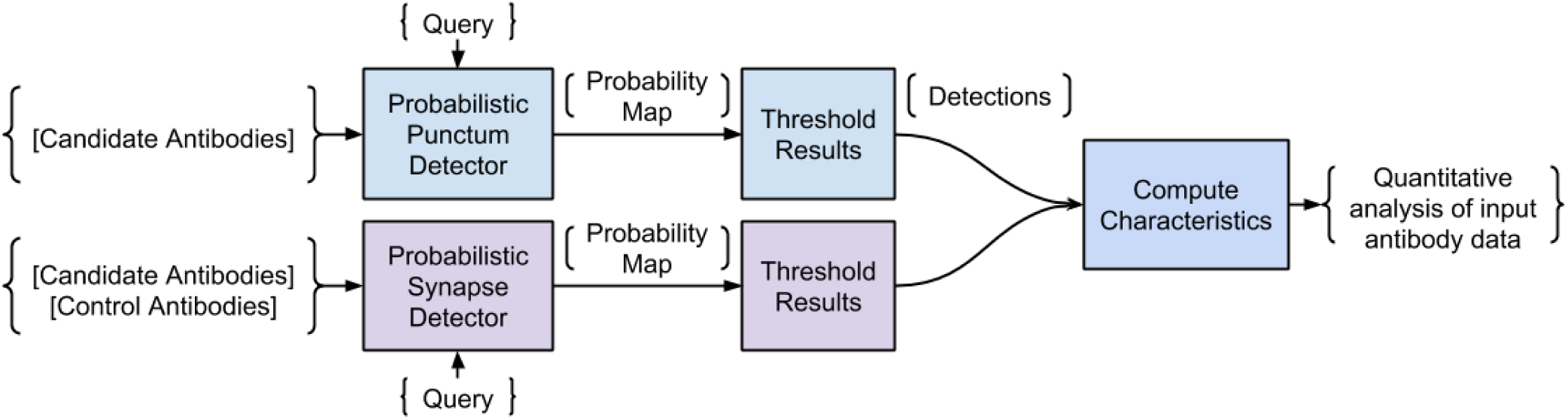
Pipeline of the Synaptic Antibody Characterization Tool (SACT). SACT combines a new probabilistic punctum detector (top row) with a probabilistic synapse detector (Simhal et al., 2017) to study the desired properties of the antibody.

The data used for validating SACT was derived from serial sections of plastic-embedded tissue that was immunofluorescently labeled with the tested antibody alongside one or more reference antibodies, chosen depending on the antigen. A selected area was then imaged on at least 3 consecutive sections, the images were aligned into stacks (Figure 3), and the performance of the tested antibody was assessed using SACT.

**Figure 3:**
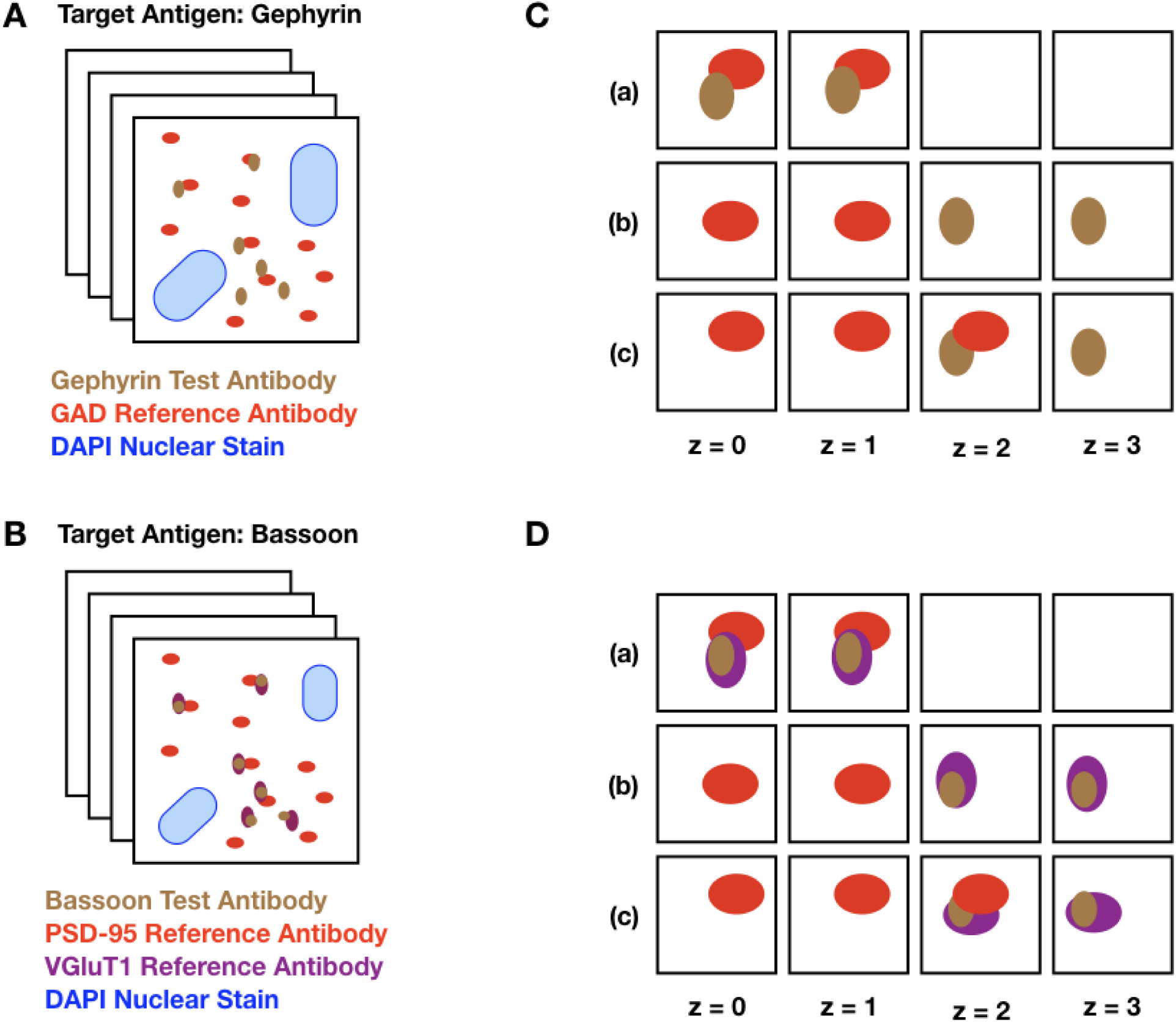
Schematic diagram of input datasets. *A.* The target protein, gephyrin, is a postsynaptic protein at inhibitory synapses, expected to be adjacent to GAD, an abundant presynaptic protein at inhibitory synapses. The set of squares represents a stack of images from serial ultrathin sections from mouse neocortex, double labeled with a gephyrin candidate antibody (brown dots) and a previously validated antibody against GAD (red dots). The large blue blobs represent DAPI, a marker for cell nuclei. *B.* Identical setup, but with Bassoon (a presynaptic protein present in excitatory synapses) as the target protein. In this example, the tissue is labeled with two reference antibodies to excitatory synapses, the presynaptic protein VGluT1 (purple dots) and the postsynaptic protein PSD-95 (red dots). *C.* Different combinations of puncta are detected on sections (z=0 to z=3) through a synapse. The test and the reference antibody can be present alongside each other on the same section (a), they can lie adjacent in the z-direction (b), or they can be adjacent both in the same section and across multiple sections (c). *D.* Identical setup as C with two reference antibodies depicted.

Importantly, SACT is applicable to a variety of synaptic antigens with very different distributions, because the user defines the expected molecular composition and size of synapses where the antigen is present. Furthermore, the algorithm can be applied to new datasets without creating extensive manual annotations for each synapse subtype, unlike traditional classifiers such as support vector machines and deep learning used by other synapse detection algorithms (Bass et al., 2017; Busse and Smith, 2013; Collman et al., 2015; Fantuzzo et al., 2017; Kreshuk et al., 2014).

### 2.2 Puncta Detection

Immunolabeling for synaptic proteins appears as small blobs or ‘puncta,’ typically less than 1 *μ*m diameter. Because synaptic structures are generally larger than the typical thickness of the individual sections used in our datasets (70 nm), the puncta corresponding to proteins that are abundant throughout the presynaptic or postsynaptic side span several sections and thus form three-dimensional puncta. Puncta detection (Figure 4) follows the general computational framework of the probabilistic synapse detector (Simhal et al., 2017), where the input is the volumetric image data and a user-defined query which includes the minimum expected 3D punctum size. Requiring a minimum 3D size minimizes the impact of random specks of noise generated during the image acquisition process and ensures that the immunolabeling is appropriately expressed across slices. For instance, a target protein that is abundantly expressed at a synapse (e.g., synapsin) should be detected across multiple slices at the current working resolution. Therefore, the presence of a punctum in only one slice likely indicates random noise, nonspecific labeling or fluorescent contaminant. On the other hand, there is little reason to assume that less abundant target proteins or those present at isolated nanodomains within synapses (e.g., many receptors or ion channels) need to span multiple slices through a synapse.

**Figure 4:**
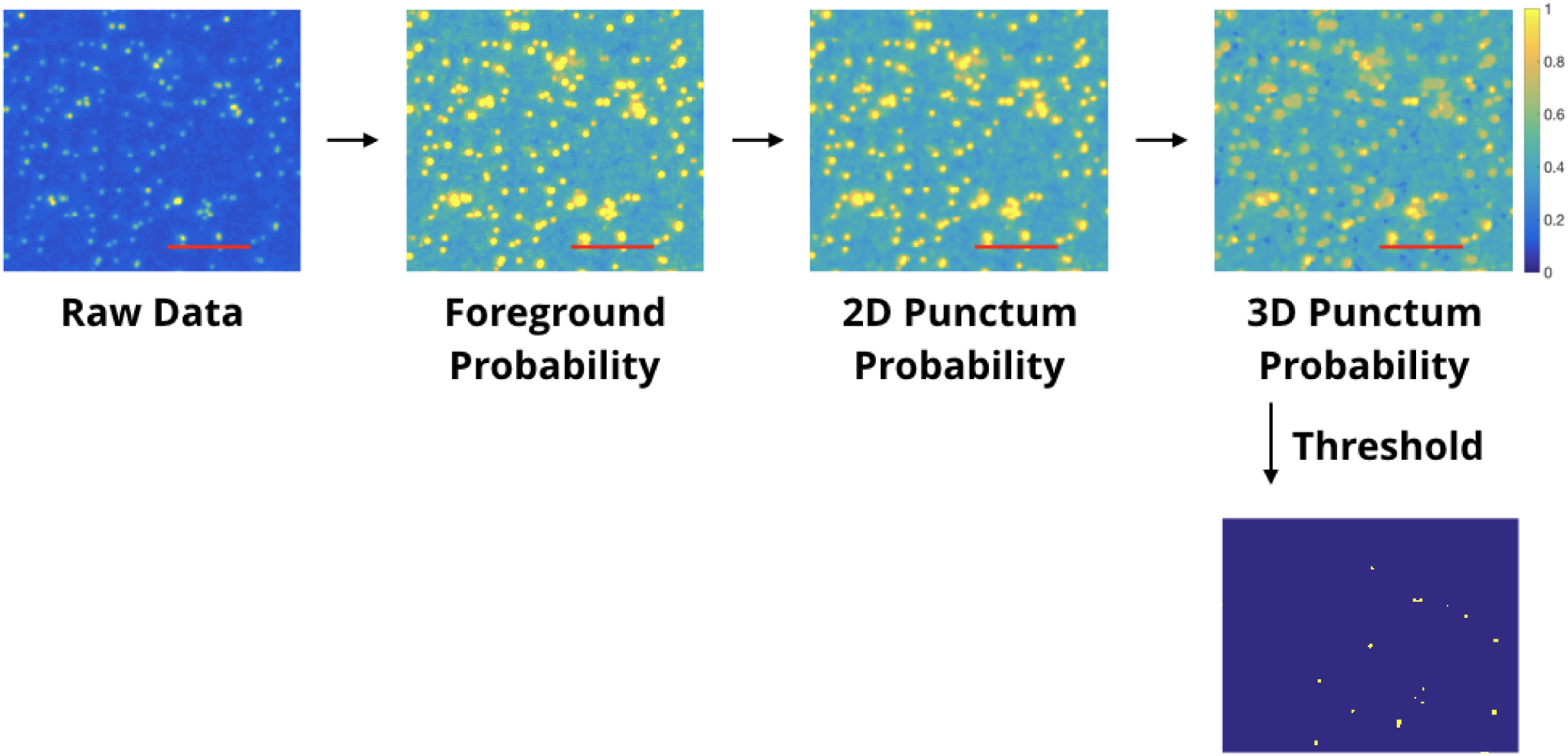
Automated punctum detection pipeline. The first box shows the raw image data from immunolabeling with one antibody. The second box is the output of a ‘foreground probability’ step; the intensity value of each pixel codes the probability it belongs to the foreground. The third box is the output of a ‘2D Punctum Probability’ step; each pixel codes the probability that it belongs to a 2D blob. The ‘3D punctum probability’ image’s intensity values are the probability that a voxel belongs to a blob which spans the minimum number of slices. The thresholded image is shown below. Red scale bars represent 5 *μ*m.

The output of the punctum detector is a probability map where the value at each voxel is the probability it belongs to a 3D punctum. To obtain the putative 3D punctum detections, this output may be thresholded if desired.

### 2.3 Synapse Detection

Characterizing synaptic antibodies for AT immunolabeling of brain sections requires detecting synapses. Over the past few years, several synapse detection methods have been presented which use traditional machine learning paradigms for detection (Bass et al., 2017; Busse and Smith, 2013; Collman et al., 2015; Fantuzzo et al., 2017; Kreshuk et al., 2014). While they perform well, each requires the user to supply manually-labeled synapse annotations for training - an often impractical and tedious requirement for antibody validation, for which manual annotations would have to be created for each antibody. The probabilistic synapse detection method introduced in Simhal et al. (2017) does not require training data for synapse detection, making it an ideal synapse detector to use for antibody characterization. The algorithm takes as input the immunofluorscence data from the candidate and reference antibodies and the expected synapse size (together referred to as the ‘query’) and outputs a probability map, where the value of each voxel represents the probability that it belongs to a synapse. This output can be thresholded in a variety of ways to obtain the putative 3D synapse detections (see (Simhal et al., 2017) for a detailed discussion). This algorithm is very flexible; as detailed below, the queries can be adapted to different data characteristics and analysis goals, further rendering it appropriate for antibody validation.

Generally, at least two known synaptic markers are required to detect a synapse, therefore, some of the datasets contain two reference antibodies to facilitate synapse detection (Figure 3B, D). However, when screening multiple candidate antibodies, it is important to minimize the time and cost of the screen. When performed with caution using appropriate antibody combinations, even a single pre-validated antibody can generate interpretable synapse-specific data. For example, if the target protein for which an antibody is being evaluated is localized at the postsynaptic sites of inhibitory synapses, as is the case with gephyrin, (Sheng and Kim, 2011), a reasonable strategy would be to label the tissue with an antibody against glutamate decarboxylase (GAD), a protein known to be specific to presynaptic terminals of GABAergic inhibitory synapses for which well-validated antibodies have already been identified. The corresponding query would then be to look for synapses containing immunolabeling for gephyrin and GAD (Figure 3A,C). If the target protein of interest is instead localized to excitatory synapses, the query may include proteins specific to excitatory synapses such as the postsynaptic protein PSD-95 (postsynaptic density 95) or the presynaptic protein VGluT1 (vesicular glutamate transporter 1). This flexibility makes this framework ideal for antibody characterization.

### 2.4 SACT Measurements

In order to evaluate and characterize antibodies, a series of measurements are computed; each measurement captures an aspect of the antibody’s performance that would be sought by an expert observer when manually interacting with the data. Each of these measurements provides unbiased quantitative information to help evaluate the intensity, specificity and sensitivity of immunolabeling obtained with a given antibody. These include the density of puncta, and the average punctum volume and standard deviation for each antibody, as well as the synapse density (number of detected synapses per volume), and target specificity ratio (the ratio of detected synapses to detected candidate antibody puncta). These measures provide the user an immediately useful quantitative assessment of the data.

#### Detected Density of Puncta

The density of the detected 3D puncta reflects the biological properties of the tissue (i.e., the abundance and distribution of the target protein), as well as the intensity and specificity of the immunolabeling with a given antibody at the concentration used. If the labeling is unexpectedly sparse, it may indicate the antibody is too highly diluted or insensitive. If it is unexpectedly dense, it may indicate that the antibody is unselective or binding to many non-synaptic sites such mitochondria or other ‘sticky’ subcellular sites. To determine the number of puncta detected in each channel, probability maps, where the value at every voxel is the probability it belongs to a 3D punctum, are computed, as described in the previous section, and then thresholded.^1^. The process is outlined in Figure 4; a value of 0 indicates no puncta were found, and the units (in this work) are in puncta per cubic micrometer. The 3D punctum density is then calculated:

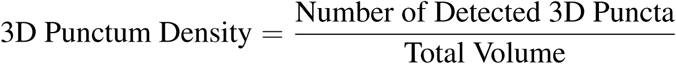

#### Average Punctum Volume

After segmenting the immunolabeling data, the average volume of puncta and their standard deviation in punctum volume are calculated. If the average punctum size is smaller or larger than expected, it may indicate a lack of efficacy or specificity for immunolabeling with that antibody. If the standard deviation of punctum volume is large, it may indicate erratic labeling. Either way, it is important to quantify the size distribution when comparing multiple antibody clones for the same target. Figure 5 shows an example of an antibody clone displaying a very large standard deviation, making it unlikely that the candidate antibody on the left will serve as a satisfactory marker for inhibitory (collybistin-positive) synapses. As before, we could compute fuzzy volumes if we prefer to work with the probability maps instead of the thresholded data. In this work, the values are in pixels.

**Figure 5:**
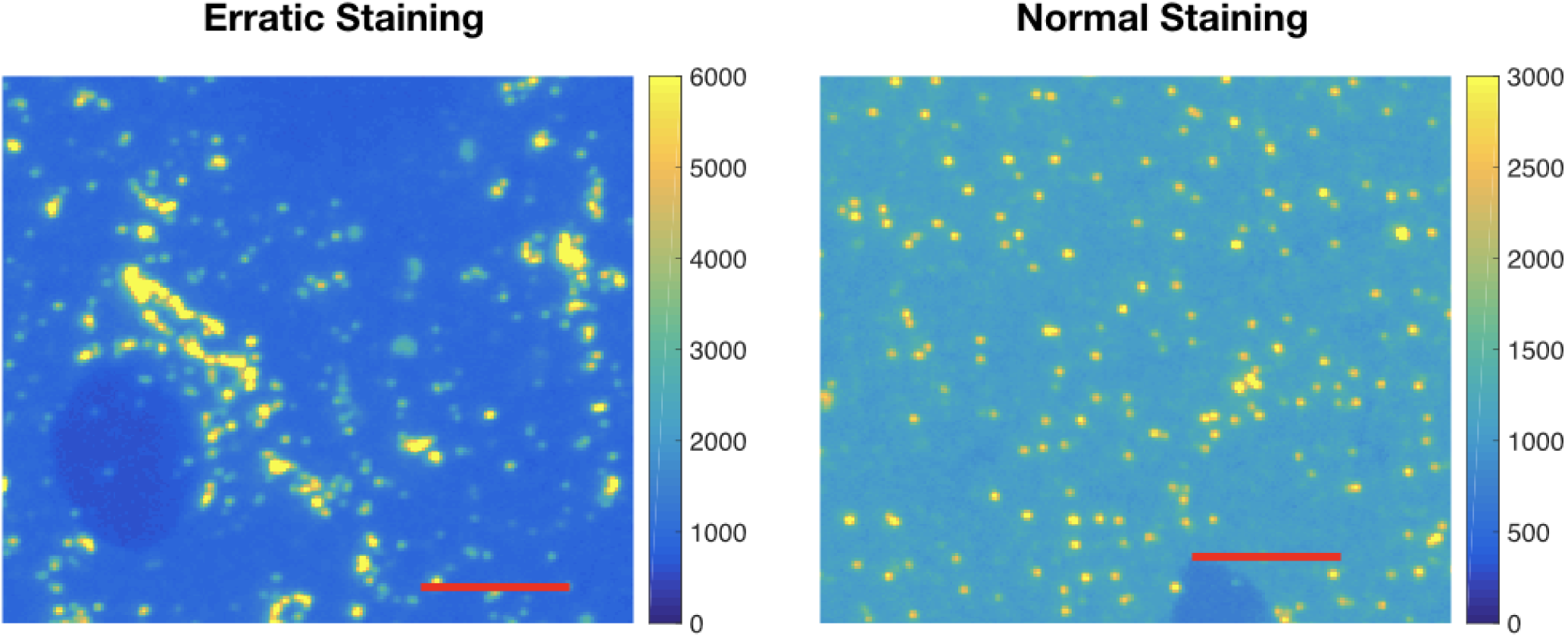
Example of erratic labeling. (*Left*) Immunolabeling for collybistin (associated with GABAergic synapses) on a raw IF slice; note the very broad distribution of sizes for puncta. (*Right*) Relatively ‘normal’ pattern of immunolabeling on a raw IF slice, using a different collybistin candidate antibody. This difference is automatically quantified by computing the average three dimensional punctum size and size variance. Each red scale bar is 5 *μ*m.

#### Synapse Density

When evaluating a synaptic antibody, it is essential to confirm that the immunolabeling localizes at the expected population of synapses. To do so, the proposed SACT incorporates a probabilistic synapse detector, the thresholded output of which is the number of synapses detected with the candidate antibody. The synapse density of a given volume is computed as

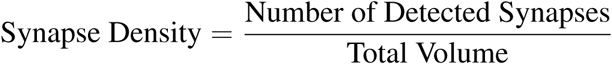

This measure is useful for evaluating antibodies against targets with a known distribution at synapses, where the density of synapses containing the target protein can be estimated. For example, ubiquitous markers of inhibitory synapses like gephyrin should be present at the large majority of inhibitory synapses, and should therefore have a synaptic density in rodent neocortex on the order of 0.15 synapses per *μm*^3^ (Knott et al., 2002). A computed synapse density substantially lower than expected may indicate low sensitivity of the antibody and/or insufficient concentration, while a synapse density considerably above that expected likely reflects nonspecific (off-target) binding of the antibody. In this work, the units used are synapses per cubic micrometer.

#### Target Specificity Ratio.

The target specificity ratio (TSR), which represents the fraction of puncta immunolabeled with the candidate antibody that are actually associated with a bona fide synapse, and is computed as

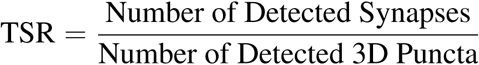

TSR ranges from 1 (every punctum detected has an associated detected synapse) to 0 (no detected punctum has an associated detected synapse); the higher the TSR, the lower is the magnitude of the nonsynaptic labeling obtained with that candidate antibody. Interpretation of this measurement will depend on the specific target; some proteins associated with synapses are almost exclusively present at synapses (e.g., synapsin), while others are also found at extrasynaptic locations (e.g., glutamate receptors). Therefore, the TSR reflects both the biological distribution of the target protein, and non-specific binding of the candidate antibody. When comparing two antibodies against the same antigen, differences in TSR reflect differences in their specificity.

## 3 MATERIALS

### 3.1 Datasets

The datasets presented were created from adult mouse neocortical tissue that was prepared, immunolabeled, and imaged using standard methods of AT (Micheva et al., 2010; Collman et al., 2015). Briefly, the tissue was chemically fixed using 4% paraformaldehyde in PBS, embedded in LR White resin, and cut into serial ultrathin sections (70 nm) that were mounted onto coverslips. One of the datasets from the automated ranking of the candidate monoclonal antibodies experiments (IRSp53) used tissue prepared in a different way: chemical fixation with 2% paraformaldehyde and 2% gluaraldehyde in PBS, followed by freeze-substitution and embedding in Lowicryl HM20 (Collman et al., 2015). The sections were labeled using indirect immunofluorescence and imaged on an automated wide-field fluorescence microscope (Zeiss AxioImager Z1, Zeiss, Oberkochen, Germany) with a 63x Plan-Apochromat 1.4 NA oil objective. While the exact size of the datasets varies, their general structure is consistent. Each dataset is composed of multichannel stacks of images from serial sections with (for the current data acquisition protocol) 100×100 nm pixel size and 70 nm slice thickness.

### 3.2 Antibodies

The antibodies used for the experiments presented here are listed in Table 1. Some of the antibodies used were tested in conjunction with screening of monoclonal antibody projects at the UC Davis/NIH NeuroMab Facility, consistent with our goal to facilitate testing of large panels of candidate antibodies in an efficient and objective fashion.

**Table 1:** Antibodies used in this study. *RRID: Research Resource Identifier. For the NeuroMab projects, the RRID of the antibody finally selected is listed; this selection was based on other factors in addition to the antibody performance evaluated using the current method.

| *Target Protein* | *Host* | *Antibody Source* | *RRID\** |
| --- | --- | --- | --- |
| Bassoon | Mouse | NeuroMab L124 project | RRID:AB_2716712 |
| Cav3.1 | Mouse | NeuroMab 75-206 | RRID:AB_2069421 |
| Cav3.1 | Rabbit | Synaptic Systems 152 503 | RRID:AB_2619850 |
| Collybistin | Mouse | NeuroMab L120 project | RRID:AB_2650452 |
| GAD2 | Rabbit | Cell Signaling 5843 | RRID:AB_10835855 |
| Gephyrin BD | Mouse | BD Biosciences 612632 | RRID:AB_399669 |
| Gephyrin | Mouse | NeuroMab L106 project | RRID:AB_2617120 |
|  |  |  | RRID:AB_2617121 |
|  |  |  | RRID:AB_2632414 |
| GluA1 | Rabbit | Millipore AB1504 | RRID:AB_2113602 |
| GluA2 | Mouse | Millipore MAB397 | RRID:AB_2113875 |
| GluA3 | Rabbit | Abcam ab40845 | RRID:AB_776310 |
| Homer1 | Mouse | NeuroMab L113 project | RRID:AB_2629418 |
| IRSp53 | Mouse | NeuroMab L117 project | RRID:AB_2619741 |
| GluN1 | Mouse | Millipore MAB363 | RRID:AB_94946 |
| PSD95 | Mouse | NeuroMab 75-028 | RRID:AB_2292909 |
| PSD95 | Rabbit | Cell Signaling 3450 | RRID:AB_2292883 |
| Synapsin | Guinea pig | Synaptic Systems 106 004 | RRID:AB_1106784 |
| Synapsin | Rabbit | Cell Signaling 5297 | RRID:AB_2616578 |
| VGAT | Mouse | NeuroMab L118 project | RRID:AB_2650550 |
| VGluT1 | Guinea pig | Millipore AB5905 | RRID:AB_2301751 |
| VGluT1 | Mouse | NeuroMab 75-066 | RRID:AB_2187693 |

## 4 RESULTS

### 4.1 Framework Evaluation

The proposed synaptic antibody characterization and screening framework was evaluated via three different tasks. Each task demonstrates an aspect of the framework necessary for validating synaptic antibodies.

#### Pairwise comparisons

Comparing the performance of two previously validated antibodies against the same synaptic target protein.

#### Concentration comparisons

Comparing the performance of a single antibody at different concentrations.

#### Evaluating candidate monoclonal antibodies

Comparing the performance of multiple antibodies raised against the same antigen.

The first and second task validate the measurements proposed in the framework and involve only antibodies previously validated for array tomography. The third task evaluates the efficacy in a ‘real world’ application-characterizing multiple antibody candidates whose suitability for AT has not yet been determined, and whose concentration is not known.

### 4.2 Pairwise Comparisons

When two well-performing antibodies are available for use in a specific application, a common question is ‘which one is better?’ Therefore, we created five AT datasets, each with two previously validated antibodies used at concentrations previously determined to yield optimal immunolabeling. These antibody pairs were evaluated alongside an antibody for a different synaptic target protein, thoroughly validated for AT in prior studies (Micheva et al., 2010; Weiler et al., 2014). An example slice of each dataset is shown in Figure 6. The higher scoring antibody was judged to be the one that had more puncta associated with the reference antibody (i.e., labels more synapses, true positives) and/or fewer puncta not associated with the reference antibody (false positives). In the dataset comparing the different PSD-95 clones, the PSD95R antibody clone had more puncta adjacent to synapsin, without noticeably more synapsin-unrelated puncta, and was therefore evaluated as better performing than PSD95M. Cav3.1R had more puncta that are not adjacent to VGluT1, and also displayed nonspecific labeling of the cell nucleus, and was therefore judged to perform worse than Cav3.1M. In the VGluT1 dataset, the difference between the two antibodies was subtle, as shown by the measurements of synaptic density and target specificity (see Table 2).

**Table 2:** Results from pairwise antibody comparisons. Punctum density, average punctum size, and standard deviation of punctum size were all computed for each antibody tested. The names in bold in the second column represent the antibodies preferred by an expert based on visual examination. In each case, the measures automatically computed by the framework agree with the expert observer’s judgement.

| Target Protein | Tested antibodies | Reference Antibody | Punctum density / $\mu\text{m}^3$ | Average punctum size / standard deviation (pixels) | Synapse Density / $\mu\text{m}^3$ | Target Specificity Ratio |
| --- | --- | --- | --- | --- | --- | --- |
| *Synapsin* | Synapsin GP | PSD-95 | 2.01 | 15.9 / 59.4 | 0.55 | 0.275 |
|  | **Synapsin R** |  | 1.09 | 27.0 / 48.6 | 0.5 | 0.457 |
| *VGluT1* | **VGluT1GP** | Synapsin | 1.54 | 12.1 / 30.2 | 0.44 | 0.283 |
|  | VGluT1M |  | 1.27 | 11.0 / 22.4 | 0.33 | 0.257 |
| *PSD-95* | PSD-95M | Synapsin | 0.95 | 10.7 / 101.7 | 0.54 | 0.568 |
|  | **PSD-95R** |  | 1.14 | 25.4 / 39.9 | 0.79 | 0.691 |
| *Gephyrin* | **GephyrinL106** | GAD | 0.72 | 12.5 / 94.1 | 0.13 | 0.181 |
|  | GephyrinBD |  | 0.46 | 17.9 / 263.1 | 0.08 | 0.175 |
| *Cav 3.1* | **Cav3.1M** | VGluT1 | 0.58 | 7.4 / 14.0 | 0.33 | 0.568 |
|  | Cav3.1R |  | 1.22 | 8.6 / 52.3 | 0.33 | 0.271 |

**Figure 6:**
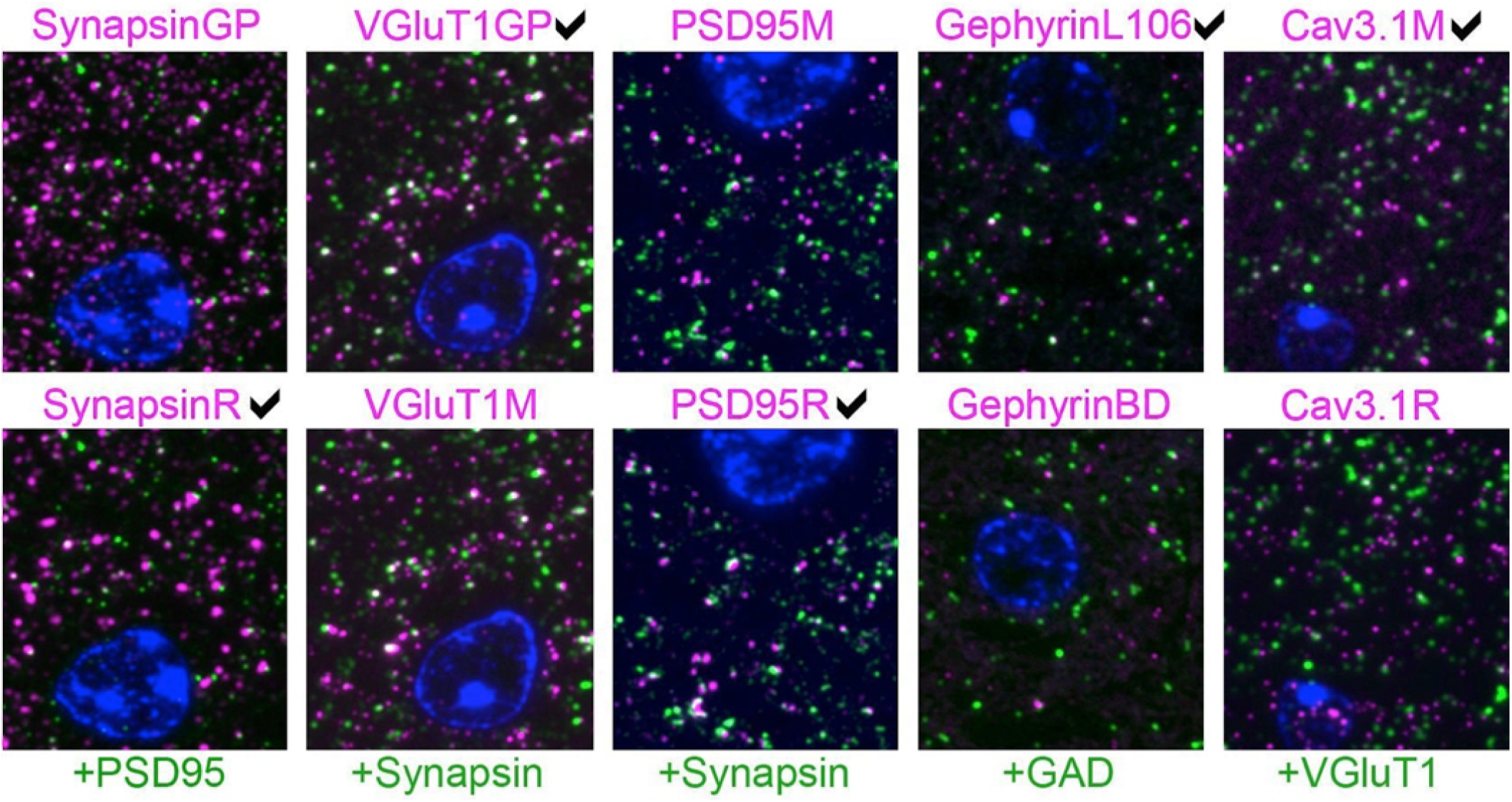
Pairwise comparison of immunofluorescence on single sections from mouse brain. Each column represents an experiment where two antibodies against the same target protein (magenta) were evaluated by double labeling with a reference antibody (green). The expert’s visually-based preference is marked for each column. The sections are also labeled with the nuclear stain DAPI (blue). For each experiment, the two images are from the same section, except for the gephyrin results, where immunolabeling with the two gephyrin antibodies was performed on different sections. Each image is 16 × 18 *μ*m.

For each dataset, the minimum expected marker size was set at 0.2 × 0.2 × 0.14 *μ*m, corresponding to 2 pixels by 2 pixels by 2 slices. Each dataset was also independently evaluated and ranked by an expert observer (KDM) blind to the automatically computed results, based on visual examination of the immunolabeling. Two measures, synapse density and target specificity ratio, were used to rank the two candidate antibodies (Table 2). When the antibodies are used at their optimal concentration, a higher measured synapse density implies higher sensitivity of the antibody (since it detects the target protein at more synapses). A higher target specificity ratio (i.e., a higher proportion of detected immunolabeled puncta that are associated with synapses) indicates higher selectivity of the antibody for the protein of interest. In each of the five cases, the antibody with the higher TSR was the same one that was visually identified by the expert as yielding the more robust and specific pattern of immunolabeling. VGluT1 and PSD-95 antibodies scored higher on both sensitivity (synapse density) and specificity (TSR), while others scored higher on only one of these measures. In the cases of Cav3.1 and synapsin, the higher-scoring antibodies had higher TSR but gave synaptic densities comparable to the other antibody. In the case of gephyrin, both antibodies had similar TSR, but the higher-ranked antibody had a higher synaptic density. These results illustrate the importance of using complementary measurements for antibody evaluation. The proposed framework provides multiple objective computations, and the user can pick the most suitable one(s) for a given task.

To evaluate the robustness of the framework, the same comparisons were performed using queries with smaller and larger minimum synapse size requirements (requiring puncta to span only one slice vs three slices), as shown in Figure 7. All queries gave consistent results for all five antibody pairs, except for Query 1, which defined a synapse as spanning only one slice; in two out of the five cases, Query 1 failed to unequivocally identify the otherwise highest scoring antibody. Thus, the use of even limited three-dimensional information from immunolabeling on serial sections as in Queries 2-4, enabled robust quantification of antibody performance.

**Figure 7:**
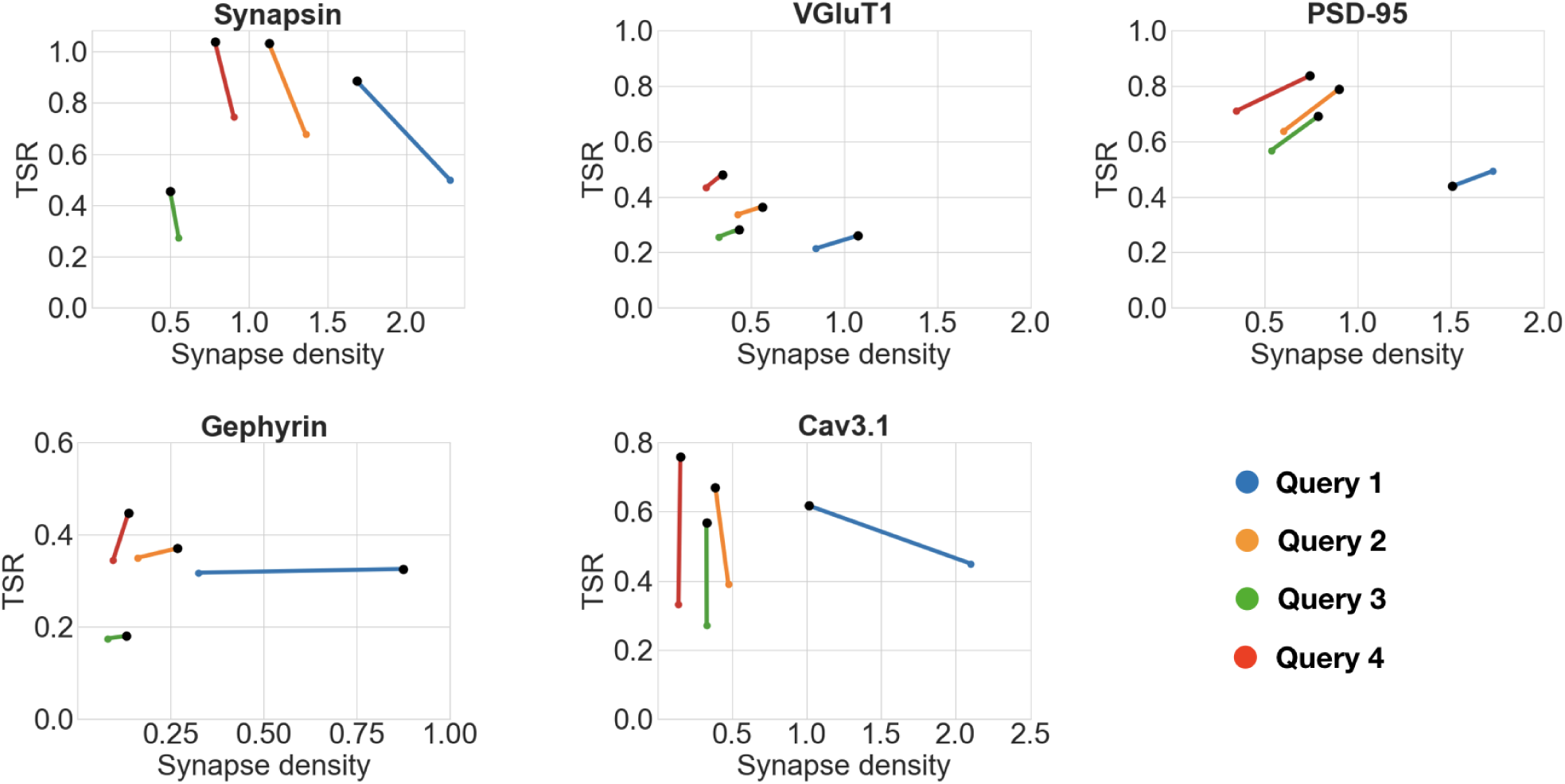
Impact of punctum size requirements on antibody comparisons. Black dots represent the higher scoring antibody. Each scatter plot shows the results of the comparison of two antibodies against the same target protein while varying the minimum synapse size requirements (queries numbered 1 through 4). Query 1 (blue): one slice of tested antibody, 1 slice control. Query 2 (orange): two slices of tested antibody, 1 slice control. Query 3 (green): two slices of tested antibody, 2 slices control. Query 4 (red): three slices of tested antibody, 1 slice control. The numerical results of query 3 are presented in Table 2. All queries gave consistent results for all five antibody pairs, with the exception of Query 1 (see text for details). Furthermore, a TSR value of greater than one (meaning more synapses than antibody puncta detected) is an artifact of thresholding dividing punctum into two, it is remedied with simple morphological operations.

This experiment illustrates the power and breadth of the proposed method. The queries can be designed by the user to take into account resolution, synapse type, and antibody binding target. Multiple queries can be run, and the antibody performance can be objectively evaluated with multiple measurements.

### 4.3 Concentration Comparisons

The optimal concentration of an antibody, which is dependent on both its binding affinity for the target protein and the abundance of the target protein in the particular sample, must be determined experimentally. Too high a concentration of the antibody will lead to high background labeling (false positives), while too low a concentration will lead to sparse labeling (false negatives). The proposed framework quantifies the effects of antibody concentration on immunolabeling of AT sections, as evaluated by the synapse density and target specificity ratio measures. The expectation is that as the antibody concentration decreases, the synapse density will also decrease.

For this experiment, datasets were generated from a series of dilutions, as shown in Table 3 and Figure 8. For each dataset, except GluN1, the minimum expected punctum size was 0.2 × 0.2 × 0.14*μ*m, corresponding to 2 pixels by 2 pixels by 2 slices. The minimum punctum size for GluN1 was (0.2 × 0.2 × 0.07 *μ*m) due to inaccuracies in the alignment of this dataset that caused inconsistencies in the positions of synapses on adjacent slices (the proposed algorithm can be easily adapted to challenges in the data, simply by changing the query).

**Table 3:** Five antibodies evaluated at different concentrations. In all experiments, a reference presynaptic antibody was included at its optimal concentration. The detected synapse density decreases as the antibody concentration decreases. For GluN1, the background noise model changed to a rayleigh distribution to better suit that specific dataset. All other datasets used the a gaussian model for the background. See (Simhal et al., 2017) for a detailed discussion.

| Target Protein | Tested Antibodies | Reference Antibody | Punctum Density/ $\mu\text{m}^3$ | Average Punctum Size / standard deviation (pixels) | Synapse Density/ $\mu\text{m}^3$ | Target Specificity Ratio |
| --- | --- | --- | --- | --- | --- | --- |
| Synapsin | SynCS 1:100 | VGluT1 | 1.23 | 41.6 / 133.6 | 0.84 | 0.68 |
|  | SynCS 1:1000 |  | 1.31 | 35.9 / 95.9 | 0.83 | 0.63 |
| GluA1 | GluR1 1:25 | VGluT1 | 0.62 | 7.9 / 51.8 | 0.12 | 0.19 |
|  | GluR1 1:125 |  | 0.43 | 9.5 / 41.4 | 0.08 | 0.18 |
|  | GluR1 1:625 |  | 0.18 | 19.3 / 180.9 | 0.05 | 0.27 |
|  | GluR1 1:3125 |  | 0.13 | 38.7 / 1117.6 | 0.03 | 0.21 |
| GluA2 | GluR2 1:25 | Synapsin | 0.89 | 10.5 / 62.3 | 0.45 | 0.51 |
|  | GluR2 1:125 |  | 0.46 | 13.9 / 29.5 | 0.30 | 0.66 |
|  | GluR2 1:625 |  | 0.27 | 15.5 / 85.8 | 0.17 | 0.64 |
|  | GluR2 1:3125 |  | 0.23 | 15.9 / 103.8 | 0.13 | 0.57 |
| GluA3 | GluR3 1:25 | VGluT1 | 1.99 | 6.5 / 11.8 | 0.42 | 0.21 |
|  | GluR3 1:125 |  | 0.81 | 8.4 / 16.4 | 0.23 | 0.29 |
|  | GluR3 1:625 |  | 0.26 | 10.8 / 23.5 | 0.07 | 0.28 |
|  | GluR3 1:3125 |  | 0.17 | 20.6 / 154.1 | 0.05 | 0.26 |
| GluN1 | GluN1 1:25 | VGluT1 | 1 | 37.2 / 1541.0 | 0.72 | 0.72 |
|  | GluN1 1:125 |  | 0.88 | 17.8 / 208.2 | 0.63 | 0.72 |
|  | GluN1 1:625 |  | 0.58 | 22.0 / 748.9 | 0.37 | 0.64 |
|  | GluN1 1:3125 |  | 0.64 | 9.8 / 54.5 | 0.39 | 0.62 |

**Figure 8:**
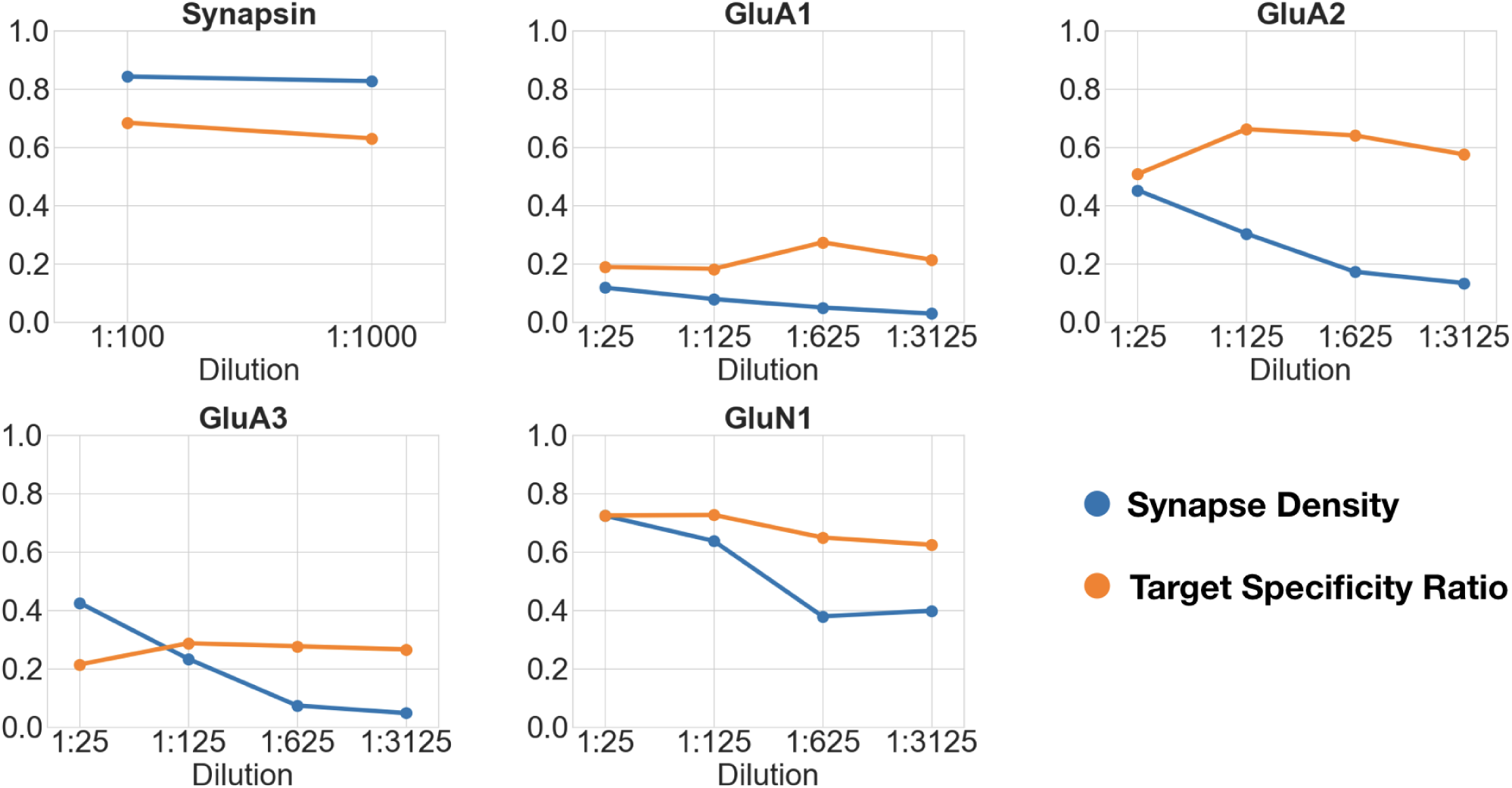
Changes in synapse density (in synapses per cubic micrometers) and target specificity ratio (TSR) as a function of antibody concentration. Each plot shows the synapse density and target specificity ratio at different concentrations of the same antibody.

The first dataset tested the effect of a 10-fold change in concentration of an antibody against the general presynaptic marker synapsin, imaged in conjunction with immunolabeling for VGluT1, a presynaptic marker of excitatory synapses. The remaining datasets tested four sequential 5-fold concentration changes on immunolabeling with different glutamate receptor antibodies, and were evaluated against immunolabeling with antibodies for general presynaptic markers (synapsin) or markers of glutamatergic synapses (VGluT1). Synapsin, previously identified as a particularly robust synaptic antibody for AT, performed equally well over the 10-fold concentration range as evaluated by both the synapse density and target specificity ratio measurements. For each of the glutamate receptor antibodies, the measured synapse density value decreased with increasing dilutions, as expected, while the target specificity value showed no consistent changes. Using the framework, we estimated that the optimal working range of the glutamate receptor antibodies is within a dilution range of 1:25 to 1:125; further dilutions led to missing too many synapses without a substantial improvement in target specificity.

### 4.4 Automated Ranking of Candidate Monoclonal Antibodies

The generation of monoclonal antibodies begins with a high-throughput screening procedure that identifies numerous candidate antibodies, all of which must then be further investigated. Because only a small fraction of these candidate antibodies will exhibit robust and specific immunoreactivity in any given application or sample preparation condition, it is important to screen as many candidate antibodies as possible for a given application. While some common antibody screens have been effectively automated (e.g., ELISA screens), screening on plastic sections from mammalian brain for antibodies that immunolabel specific populations of synapses must still be performed and analyzed manually by an expert observer, a difficult and labor-intensive process (Gong et al., 2016). The framework proposed here might facilitate the analysis of large-scale screens on tissue sections. Accordingly, we tested its performance by screening candidate monoclonal antibodies against synaptic target proteins generated at the UC Davis/NIH NeuroMab facility. This procedure is more challenging because the concentration of antibody in hybridoma tissue culture supernatants is unknown, so immunolabeling must be performed at antibody concentrations that may differ for different candidate antibodies, and these concentrations may not be optimal.

We imaged AT sections immunolabeled with a set of candidate antibodies against the same target protein. For each dataset, tissue from mouse neocortex was prepared using standard AT methods and had at least two antibodies applied - the candidate antibody for the target protein of interest, and a validated antibody at its optimal concentration. The ranking of candidate antibodies was determined based on two measurements provided by the framework: synapse density and the target specificity ratio. The target specificity ratio was the deciding factor in most cases; synapse density was used to exclude candidate antibodies with unreasonably high values based on previous biological knowledge: excitatory synapses are expected to have a density of approximately 1 per *μm*^3^, and the inhibitory synapses approximately 0.15 per *μm*^3^ (Calverley and Jones, 1987; Schüz and Palm, 1989; Knott et al., 2002). Each dataset was blindly evaluated and ranked by an expert observer, based on visual examination of the images. Screening of the Bassoon candidate antibodies was performed in two rounds; the second round included only those candidates identified as best or unclear in the first round. Overall, these experiments addressed several questions: 1) Can the framework be used to correctly rank the performance of multiple candidate antibodies?; 2) What is the minimum number of reference antibodies required to accomplish this?; and 3) What is the optimal minimum puncta size needed? In total, six datasets, ranging from 4 to 19 different candidate antibodies each, were analyzed.

#### Can the framework correctly rank the performance of multiple candidate antibodies?

Analysis of the six datasets demonstrated an excellent correspondence between the framework’s ranking and expert evaluation of candidate antibody performance. In all but one case, the framework identified all of the expert-chosen antibodies, and it only missed one in the case of gephyrin, because the group of failed candidate antibodies was too small to serve as an adequate comparison. The results are summarized in Table 4 and Figure 9.

**Table 4:** Summary of candidate antibody comparisons. Multiple candidate antibodies against the target protein were ranked using measurements from the proposed SACT. The candidate antibodies were independently evaluated by visual inspection of the immunofluorescence images. For each target protein, the reference antibodies were chosen according to the known characteristics of synapses expressing the target protein.

| Target Protein | Reference antibodies | Total # candidate antibodies tested | # good candidates chosen by expert | # good candidates chosen by SACT | SACT false negatives | SACT false positives |
| --- | --- | --- | --- | --- | --- | --- |
| Gephyrin | GAD | 7 | 5 | 4 | 1 | 0 |
| Homer | PSD-95 | 4 | 1 | 1 | 0 | 0 |
| IRSP53 | PSD-95, VGluT1 | 12 | 2 | 2 | 0 | 0 (+1 unclear) |
| VGAT | GAD | 17 | 4 | 4 | 0 | 0 (+2 unclear) |
| Collybistin | GAD | 19 | 6 | 6 | 0 | 1 |
| Bassoon 1st exp. | Synapsin | 19 | 4 | 4 | 0 | 0 (+1 unclear) |
| Bassoon 2nd exp. | VGluT1, PSD-95 | 10 | 5 | 5 | 0 | 0 |

**Figure 9:**
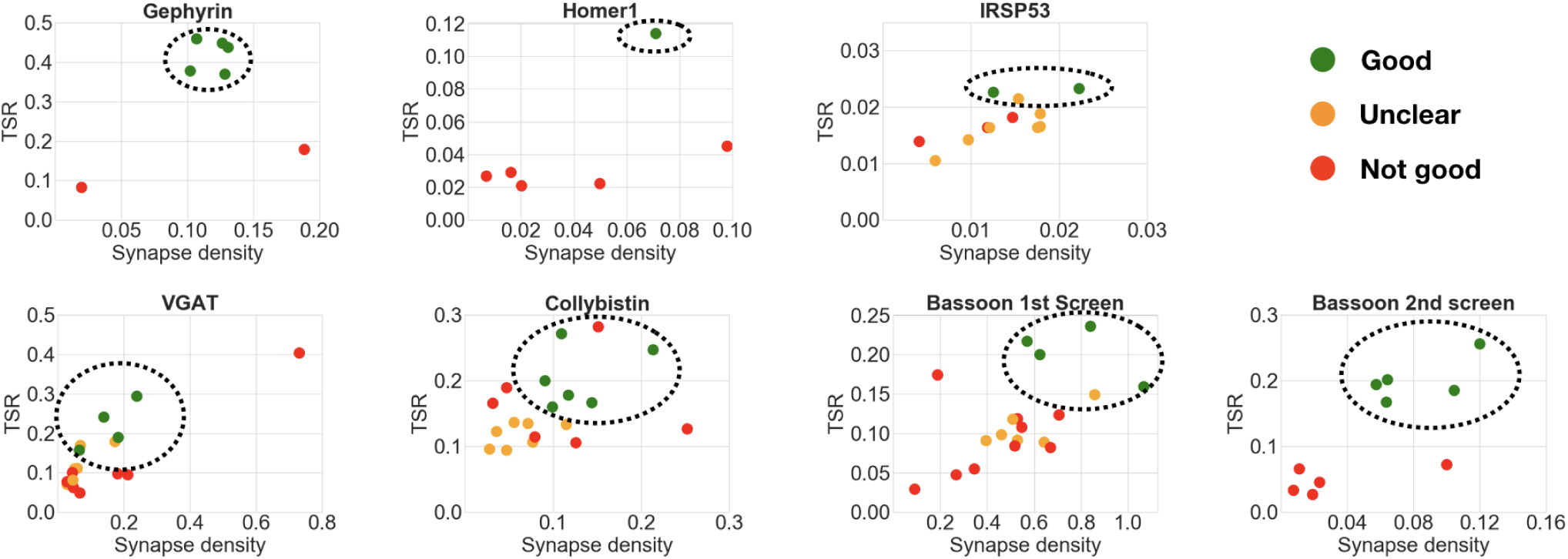
Comparison of multiple candidate antibodies. Each scatter plot shows the computed synapse density and target specificity ratio of multiple candidate antibodies, with the best-ranking candidates circled. Expert ranking is color-coded: green - best, orange - unclear, red - fail. The specific query in each case is indicated in the plot title, where the number indicates the minimum punctum size for each label. For example, ‘2 Gephyrin, 2 GAD’ means that a synapse in this case is defined as a gephyrin punctum that spans at least 2 slices adjacent to a GAD punctum that also spans at least 2 slices. The outlier in the VGAT scatter plot was not included in the best candidate antibodies selection, because of the abnormally high synaptic density (0.7 per *μm*^3^ compared to target max density of 0.15 per *μm*^3^). Screening of the Bassoon project was performed in 2 rounds: the candidate antibodies identified as best or unclear in the first round were screened again with adjusted concentrations.

#### What is the minimum number of reference antibodies?

The previous experiments with pairwise or concentration comparisons were performed using only one reference antibody. In those cases, the tested antibodies were already known to recognize their target in plastic sections; it is therefore reasonable to assume that the combination of one reference synaptic antibody with one tested synaptic antibody will generate synapse-specific data. To verify whether an additional reference antibody may offer an advantage when screening new antibodies, two of the datasets included both presynaptic and postsynaptic reference antibodies. In these two datasets, the performance of the query containing an additional reference antibody was compared to the standard single reference antibody query used in the previous experiments. In both cases, the results of the two queries were very similar (compare Figure 10 with Figure 9), suggesting that inclusion of a second reference synaptic antibody is unnecessary for the purpose of screening large sets of candidate antibodies.

**Figure 10:**
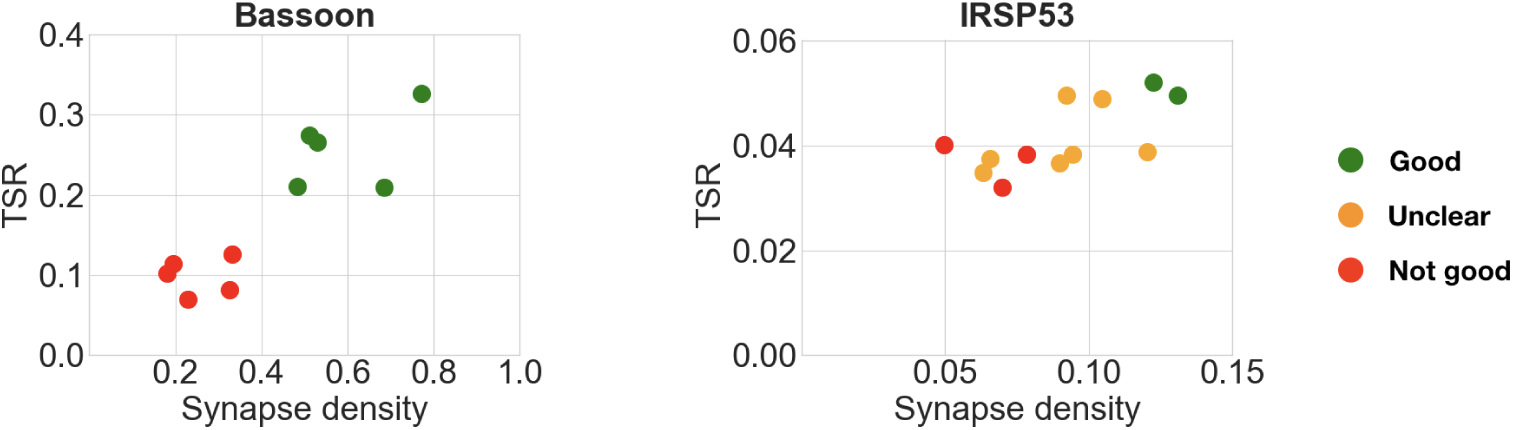
Comparison of multiple candidate antibodies using two reference synaptic antibodies. The specific query in each case is indicated in the plot title. For example IRSp53, PSD, VGluT1 means that a synapse is defined as the colocalization of an IRSp53, PSD-95 and VGluT1 punctum. Expert ranking is color coded: green - best, orange - unclear, red - fail. Compare with Figure 9.

#### What is the optimal minimum punctum size?

The pairwise antibody comparison experiments showed that the results were not affected by the stringency of the query, except in cases when the minimum puncta requirements were too permissive (smallest synapse size: labels present on 1 slice). Therefore, to screen multiple candidate antibodies, we generally chose queries of medium stringency, requiring the labels to be present in two consecutive slices. This strategy worked very well for candidate antibodies directed against abundant synaptic proteins (gephyrin, Homer1, IRSp53, VGAT, collybistin). In contrast, the permissive query, which required the labels to be present on only one section, gave inconclusive results in most of these cases (Figure 11). The first round of screening for Bassoon antibodies was an exception, because it yielded clearer results with a one-section query. This is likely due to the wide variations in concentration of the candidate antibodies present in the tissue culture supernatants used for screening, many of which required subsequent dilution, as performed in the second round of testing. In this second round with adjusted concentrations, the two-section query performed well, as seen for the other abundant synaptic target proteins. These experiments suggest that it is best to start an antibody evaluation using a query that requires the labels to be present in two slices. The top-ranking antibodies based on such a query can then be selected and visually examined by experts to confirm their performance.

**Figure 11:**
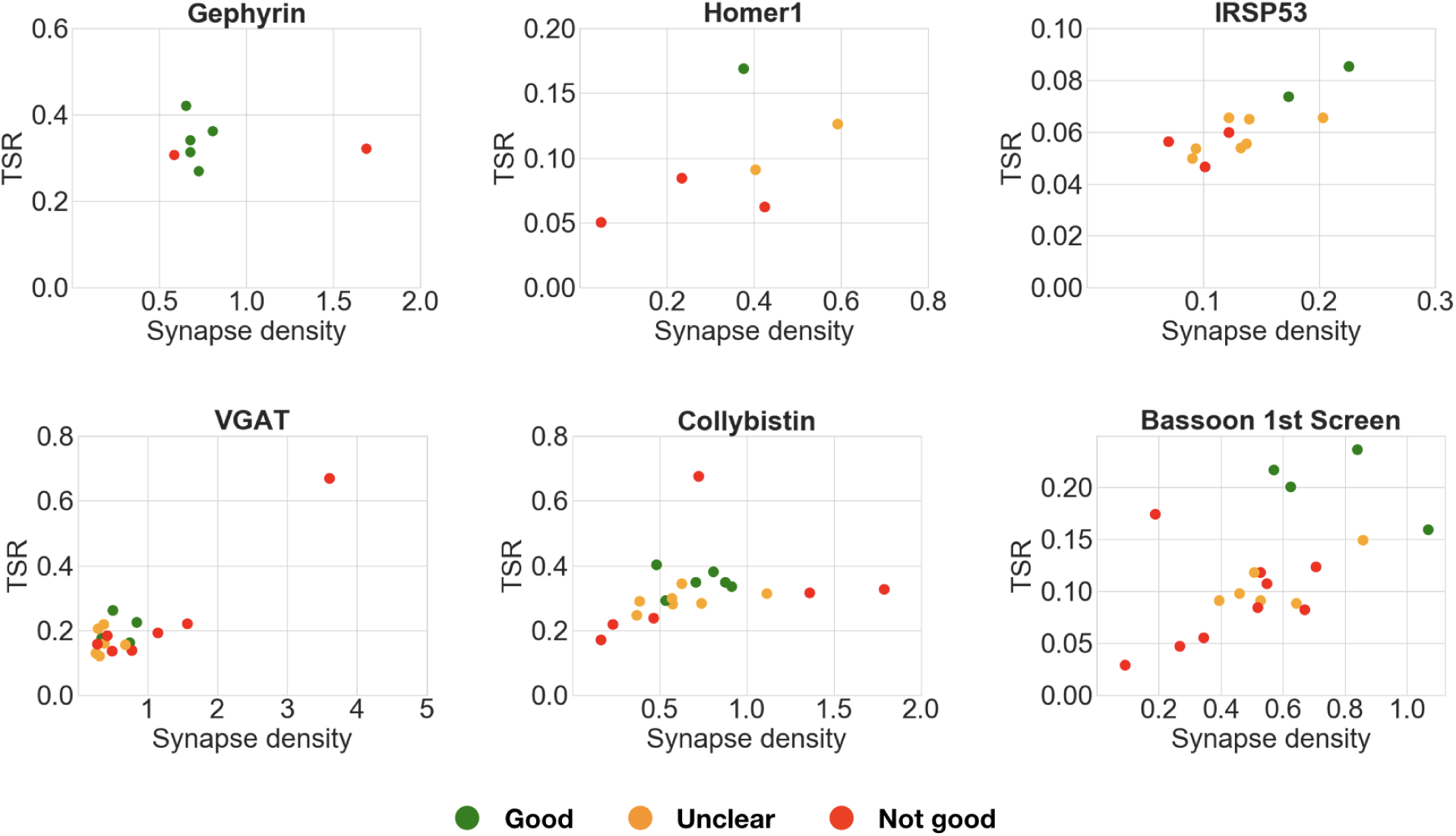
Comparison of multiple candidate antibodies and queries. These plots compare multiple candidate antibodies using an alternative query requiring the puncta to be present on only 1 slice, instead of 2 as in Figure 9. Each scatter plot shows the synapse density and target specificity ratio of multiple candidate antibodies against the same target protein. Expert ranking is color coded: green - best, orange - unclear, red - fail. The specific query in each case is indicated in the plot title, where the number indicates the minimum punctum size for each label. In many of these cases (gephyrin, VGAT, collybistin), the more permissive 1-slice query does not allow correct selection of the best performing candidate antibodies.

## 5 DISCUSSION

The present report introduces an effective framework for automated characterization and screening of antibodies for AT. The framework provides a number of automatically computed characteristics, such as synapse density and target specificity ratio, that reflect the sensitivity and specificity of immunolabeling with a given antibody. Taken together, these computed characteristics provide an objective way to characterize and compare the performance of different antibodies against the same target, simplifying the process for selecting antibodies best suited for AT. When evaluating multiple candidate antibodies, this represents an efficient method to identify a small number of promising antibodies for further evaluation.

The Synaptic Antibody Characterization Tool (SACT, the implementation of the framework) is designed to be a flexible tool for antibody screening. Because it is query-based, it allows the user to define the molecular composition and size of synapses expected to contain the antigen. This flexibility is advantageous for synaptic antibody screening because the query can be designed to focus on different synapse subtypes, depending on prior biological knowledge (e.g., what combinations of proteins are likely to be present, and where the antibody target is expected to be located). Its inherent flexibility should allow this approach to be used also to validate antibodies that target other subcellular structures, ranging from the nodes of Ranvier on myelinated axons, to mitochondria, to histone marks in the nucleus. The method works with a wide selection of reference antibodies, which need not colocalize with the tested antibody. For example, antibodies to gephyrin and collybistin, both postsynaptic proteins, were evaluated using the presynaptic marker GAD as reference. The flexibility in reference antibody selection enables users to optimize the use of their available antibody stocks. With further practical experience we anticipate that a restricted group of well-characterized antibodies will be adopted as controls for each target category.

Our experiments demonstrate that SACT provides a robust method for antibody screening, ranking antibodies based on quantitative measures of their performance. In the pairwise comparisons of antibodies, there was 100% agreement between the expert ranking and the automated antibody ranking based on synapse density and target selectivity ratio. Variations in the size requirement did not affect the ranking, as long as synapse detection was based on more than one slice. Even when a synapse was required to be present on only one slice, performance was only modestly degraded, such that the outcome measures for some antibody pairs were ambiguous. The present approach also accommodates variations in antibody concentration, as demonstrated by the experiments with multiple candidate antibodies from monoclonal antibody projects, which showed a high correlation between the ranking by algorithm and by expert evaluation of candidate antibody performance in all six datasets, even though the concentration of antibodies in the hybridoma tissue culture supernatants used for screening was unknown and intrinsically variable. This insensitivity to antibody concentration is very important in practice when evaluating multiple antibodies; by eliminating the need for immunolabeling with series of antibody dilutions, it substantially reduces the amount of work involved.

There are some limitations to the use of the proposed framework for antibody validation. This is not a stand-alone tool for generic antibody validation; it is designed to specifically address the performance of the antibody for immunofluorescence AT, and must be used along with other tests and controls. For example, SACT does not test for cross-reactivity with other proteins. A second limitation is that this approach requires prior knowledge of the expected distribution of the antigen (or some other characteristic to use for reference), especially if it is found only in a small population of synapses. In such cases it will be important to ensure that the tissue sample used for immunolabeling contains such synapses at a reasonable density and/or includes a reference marker to independently identify this population.

A number of technical issues can interfere with performance. Proper alignment of the sections in the imaged series is required to ensure that position of synapses is consistent on adjacent sections. In one of the concentration comparison experiments with GluN1, inaccuracies in the alignment led to poor performance of the algorithm when using the standard size requirement of a synapse to be present on at least two consecutive sections. In this case, a one-slice size requirement was successfully used, but we show that this approach will not always work. To fully benefit from the advantages of using three-dimensional information from multiple serial slices, one must ensure that the datasets are well aligned. Another technical issue to consider is possible bleed-through during the fluorescent imaging, which can cause the false impression of colocalization between the tested and reference antibodies.

When carefully planned and executed to avoid pitfalls, the automated framework described here can be used to identify and characterize antibodies against a wide assortment of synaptic target proteins that yield robust and specific immunolabeling in plastic sections of brain tissue. This is particularly important because despite extensive work, the nature of synaptic processing is still poorly understood, and many basic questions remain. For example, how many different types of synapse exist? How do these different types vary over different brain areas? How does their distribution change over time? With experience? Under pathological conditions? For questions of this nature, it is important to objectively assess a large number of individual synapses, and a large number of different molecules at each synapse, as can be done using AT. Identifying reliable synaptic antibodies for AT will remove a major limitation for such studies and allow a better understanding of synapses.

## ETHICS STATEMENT

All procedures related to the care and treatment of animals were approved by the Administrative Panel on Laboratory Animal Care at Stanford University.

## AUTHOR CONTRIBUTIONS

AKS, RJW, GS, and KDM designed the study, with KDM leading the effort. AKS and GS designed the computational algorithm. AKS wrote the code and ran the tests. KDM provided the data and evaluation. BG and JST provided the tested antibodies. SJS oversees the effort. AKS, RJW, GS, and KDM wrote the core of the paper with contributions from all authors.

## ACKNOWLEDGMENTS & FUNDING

This work was supported by the National Institutes of Health (NIH-TRA 1R01NS092474, R01NS094499, R01MH111768, R24NS092991, and R01NS039444.) and the Allen Institute for Brain Sciences (AIBS), along with the National Science Foundation (NSF), United States Office of Naval Research (ONR), United States Army Research Office (ARO), and the National Geospatial-Intelligence Agency (NGA).

## DATA AVAILABILITY STATEMENT

The datasets generated for this study can be found at https://github.com/aksimhal/SynapseAnalysis.

## Footnotes

1 The threshold could be avoided by changing the formula to be the sum of the probabilities instead of the thresholded number of 3D puncta. This applies to the other measurements described next as well.

